# Evolution of human, chicken, alligator, frog and zebrafish mineralocorticoid receptors: Allosteric influence on steroid specificity

**DOI:** 10.1101/151233

**Authors:** Yoshinao Katsu, Kaori Oka, Michael E. Baker

**Affiliations:** Graduate School of Life Science, Hokkaido University, Sapporo, Japan.; Department of Biological Sciences, Hokkaido University, Sapporo, Japan.; Division of Nephrology-Hypertension, Department of Medicine, University of California, San Diego, CA, USA.

**Keywords:** aldosterone, corticosteroids, progesterone, spironolactone, vertebrate MR, zebrafish, mineralocorticoid receptor, mineralocorticoid, evolution

## Abstract

We studied the response to aldosterone, 11-deoxycorticosterone, 11-deoxycortisol, cortisol, corticosterone, progesterone, 19-norprogesterone and spironolactone of human, chicken, alligator, frog and zebrafish full-length mineralocorticoid receptors (MRs) and truncated MRs, lacking the N-terminal domain (NTD) and DNA-binding domain (DBD), in which the hinge domain and ligand binding domain (LBD) were fused to a GAL4-DBD. Compared to full-length MRs, some vertebrate MRs required higher steroid concentrations to activate GAL4-DBD-MR-hinge/LBD constructs. For example, 11-deoxycortisol activated all full-length vertebrate MRs, but did not activate truncated terrestrial vertebrate MRs and was an agonist for truncated zebrafish MR. Progesterone, 19-norProgesterone and spironolactone did not activate full-length and truncated human, alligator and frog MRs. However, at 10 nM, these steroids activated full-length chicken and zebrafish MRs; at 100 nM, these steroids had little activity for truncated chicken MRs, while retaining activity for truncated zebrafish MRs, evidence that regulation of progestin activation of chicken MR resides in NTD/DBD and of zebrafish MR in hinge-LBD. Zebrafish and chicken MRs contain a serine corresponding to Ser810 in human MR, required for its antagonism by progesterone, suggesting novel regulation of progestin activation of chicken and zebrafish MRs. Progesterone may be a physiological activator of chicken and zebrafish MRs.

## Introduction

The mineralocorticoid receptor (MR) belongs to the nuclear receptor family, a diverse group of transcription factors that also includes receptors for androgens (AR), estrogens (ER), glucocorticoids (GR) and progestins (PR), as well as other small lipophilic ligands, such as thyroid hormone and retinoids (*1-4*).

Aldosterone (Aldo) is the physiological activator for human MR in epithelial tissues, such as the kidney distal collecting tubules and the colon (*5-9*). The human MR has similar strong binding affinities for several corticosteroids: Aldo, cortisol (F), corticosterone (B) and 11-deoxycorticosterone (DOC), and for progesterone (Prog) (*10-12*) (Fig. 1). These steroids also have similar affinities for rat MR (*13-15*) and guinea pig MR (*14, 15*). Corticosteroids are transcriptional activators of human MR (*10, 12, 16-18*), while, in contrast, Prog is an antagonist for human MR (*12, 17-19*) (Fig. 1). Complicating Aldo activation of human, rat and mouse MRs is the substantially higher concentration in human serum of F and in rat and mouse serum of B compared to Aldo. For example, the concentration of F in human serum is from 500 to 1,000 times higher than that of Aldo, and under stress F increases further. In this case, human MR would be expected to be occupied by F, to the exclusion of Aldo (*5, 8, 20-22*). One contributor to selective occupation of the MR in epithelial cells by Aldo over F and B arises from 11β-hydroxysteroid dehydrogenase-type 2 (11β-HSD2), which selectively inactivates F and B (*5, 22-25*). Aldo is inert to 11β-HSD2, as is DOC, which lacks an 11β-hydroxyl, allowing both steroids to activate the MR in epithelial tissues. The MR also is found in brain, brain, heart, aorta, lung, adipose tissue, breast and ovary (*6, 7, 9, 26*), some of which lack 11β-HSD2. In those tissues, F and B would be expected to occupy the MR. Also important for selective occupation of the MR by Aldo is corticosteroid binding globulin, which preferentially sequesters F, B and DOC compared to Aldo (*13, 27*). CBG has not been isolated from fish, leaving the role serum proteins in corticosteroid action uncharacterized.

**Figure 1.**
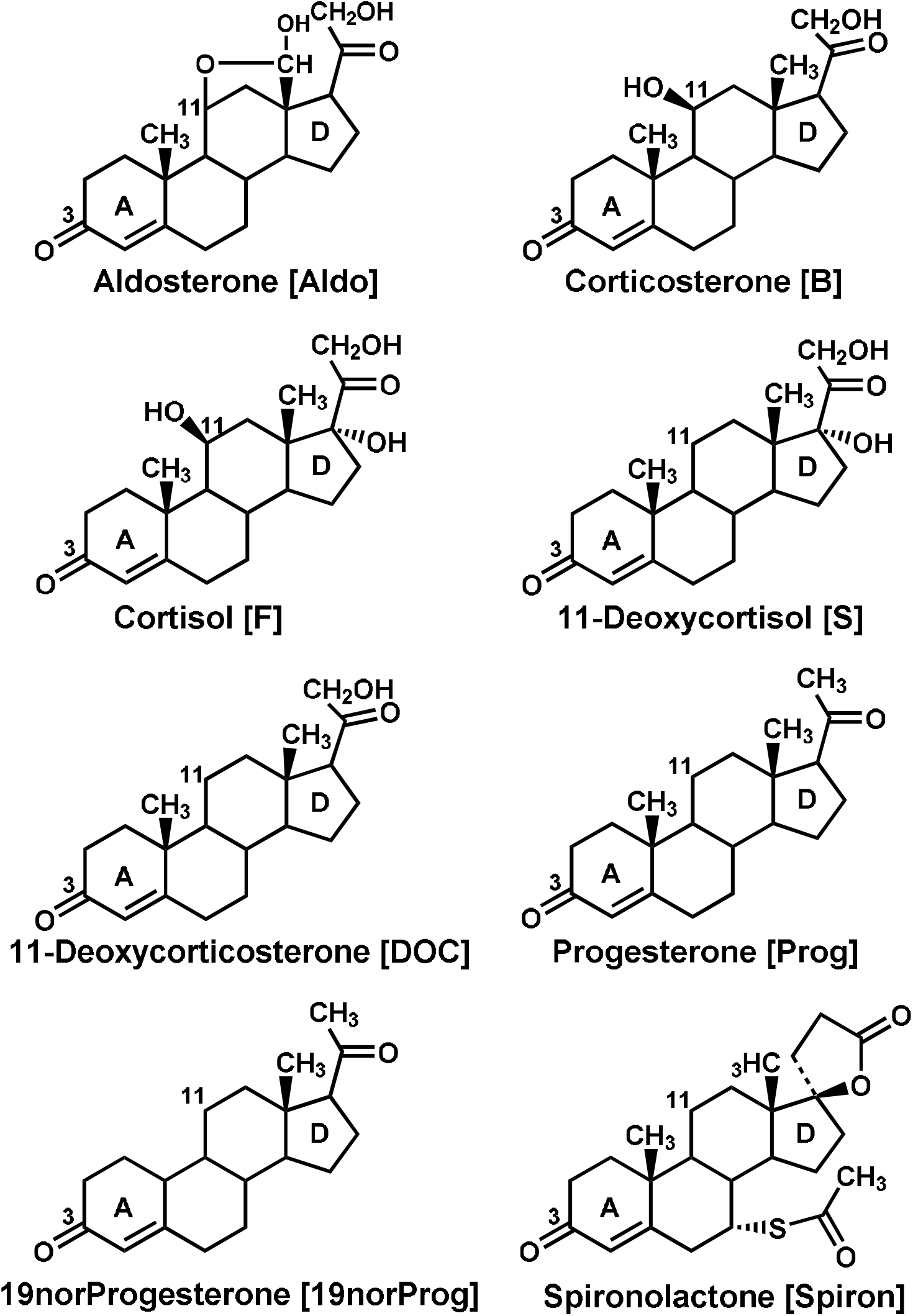
Corticosteroid and Progestin Structures. Aldosterone (Aldo) the physiological ligand for terrestrial vertebrate MRs. Cortisol (F), corticosterone (B), 11-deoxycorticosterone (DOC) also are ligands for terrestrial vertebrate MRs. 11-deoxycorticosterone (DOC) and Cortisol (F) have been proposed to be mineralocorticoids in teleosts (*18, 53-55*) because Aldo is not found in fish (*56*). Prog also may be a mineralocorticoid in fish (*11, 18, 34*). 11-deoxycortisol (S) is a ligand for corticosteroid receptor (CR) in lamprey (*57, 58*). Prog has high affinity for human MR (*12, 17, 18*), but is an antagonist at 10 nM. Spiron is an MR antagonist. However, Prog, 19norProg and Spiron are agonists for gar, sturgeon and zebrafish and trout MRs (*18, 30, 34*).

Despite the similar binding affinities of Aldo, F, B and DOC for the human MR, there is substantial variation in the half-maximal response (EC50) among these steroids for transcriptional activation of the MR. For example, Aldo has a substantially lower EC50 (higher activity) than F for human MR (*12, 16-18, 28-30*). Also, fish MRs have a stronger response to Aldo than to F, B and S (*18, 30-34*). The basis for this difference among corticosteroids in transcriptional activation of these vertebrate MRs is still not well understood.

Interestingly, Prog, 19nor-progesterone (19norProg) and spironolactone (Spiron) (Fig. 1), which are antagonists for human MR (*12, 17, 18*), are agonists for several fish MRs (*18, 30, 34*) (new Fig. 2). Data for progestin activation of frog, alligator and chicken MRs are absent. Thus, the timing of the evolution of antagonist activity of progestins and Spiron for the MR is not known.

**Figure 2.**
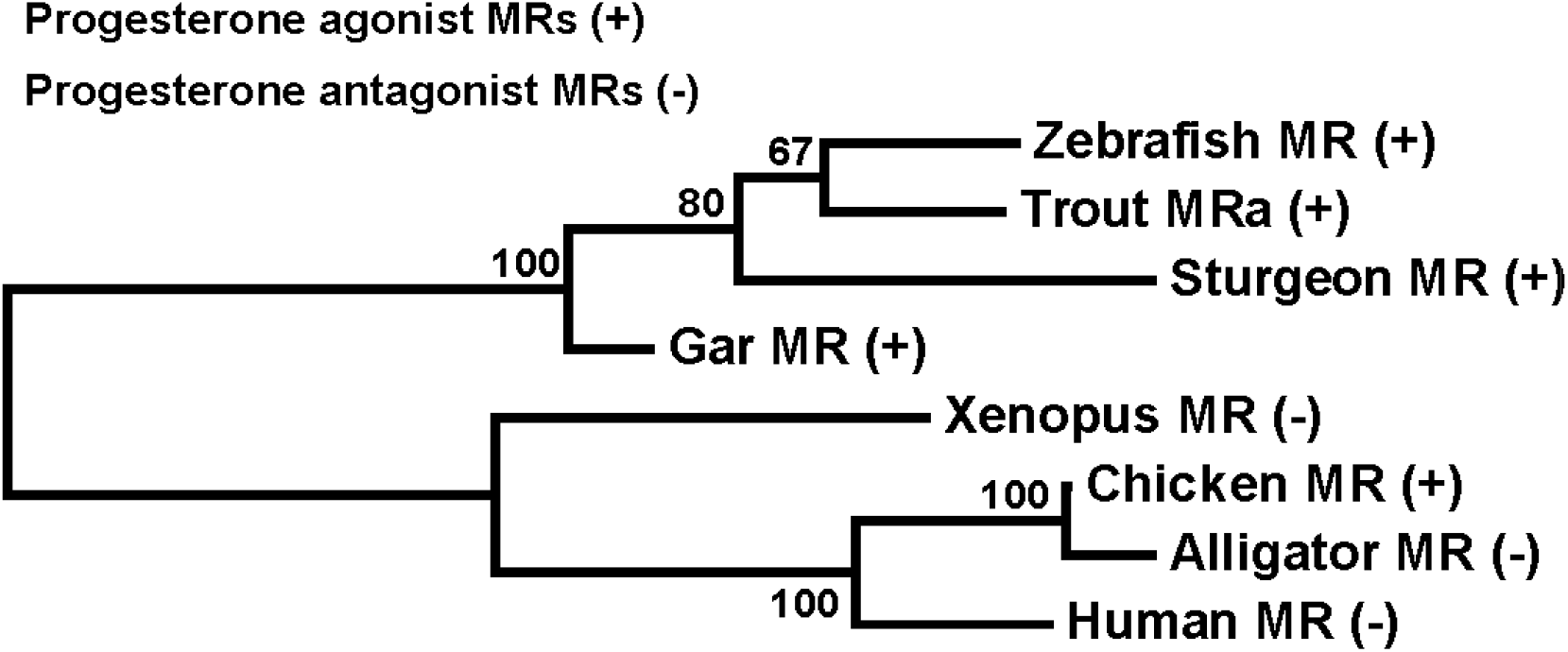
Phylogeny of vertebrates investigated for MR activation by progesterone. The ligand-binding domains of the MR from fish and terrestrial vertebrates that have been tested for activation by progesterone were collected and a phylogeny constructed with the Neighbor-Joining Method (*59*) after sequences were aligned by ClustalW (*60*). Values for 1,000 bootstrap runs are shown as percentages at each node. MR activation by Prog has been found for gar, sturgeon and zebrafish and trout MRs (*18, 30, 34*) and for chicken MR (this study). GenBank accession numbers: human MR (NP_000892), chicken (ACO37437), alligator MR (NP_001274242), *Xenopus* MR (NP_001084074), zebrafish MR (NP_001093873), trout MRa (NP_001117955), sturgeon MR (BAV17690), gar MR(BAV17691).

An important structural property that influences transcriptional activation of the MR and other steroid receptors is their modular domain structure, which consists of an N-terminal domain (NTD) (domains A and B), a central DNA-binding domain (DBD) (domain C), a hinge domain (D) and a C-terminal ligand-binding domain (LBD) (domain E) (*18, 29, 35-37*) (Fig. 3). Although the LBD alone on the MR is competent to bind steroids (*20, 35, 38-41*) allosteric interactions between the LBD and NTD are important in transcriptional activation of the human and zebrafish MR (*19, 30, 37, 42*)), as well as for the GR and other steroid receptors (*29, 43-52*).

**Figure 3.**
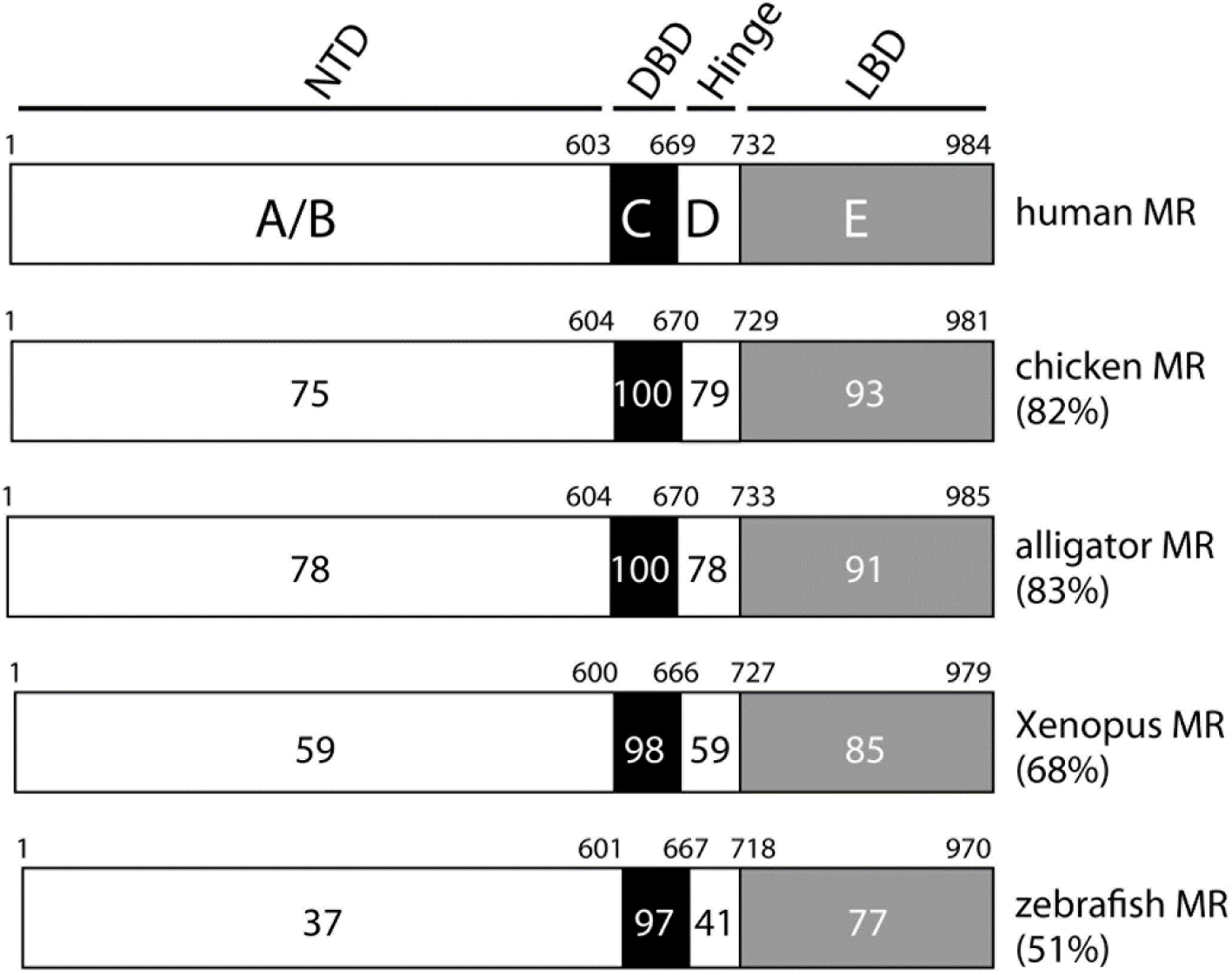
Comparison of domains in vertebrate MRs. Domains A/B (NTD), C (DBD) D (hinge) and E (LBD) on MRs from human, chicken, alligator *Xenopus* and zebrafish are compared. Shown are the number of amino acids in each domain and the percent identical amino acids compared to human MR. GenBank accession numbers: human MR (NP_000892), chicken (ACO37437), alligator MR (NP_001274242), *Xenopus* MR (NP_001084074), zebrafish MR (NP_001093873).

Moreover, there are differences between the effects F and DOC on transcription due to interactions between the LBD and NTD in human MR (*19, 37*) and zebrafish MR (*30*). In human MR, DOC and F weakly promote the NTD/LBD interaction and gene transcription (*19*). In contrast, in zebrafish MR, F and DOC substantially induce the NTD/LBD interaction and increase transcription. The basis for these differences between human and zebrafish MR is not known, as well as the effect, if any, of inter-domain interactions on corticosteroid and progestin-mediated transcription in frog, alligator and chicken MRs.

To begin to fill in these gaps, we investigated activation of full-length MRs from human, chicken, alligator, frog (*Xenopus laevis*) and zebrafish and their truncated MRs, consisting of the GAL4 DBD fused to the D domain and E domain of the MR (MR-LBD), by a panel of corticosteroids (Aldo, F, B, DOC, 11deoxycortisol (S)) and progestins (Prog, 19norProg) and Spiron. We found substantial differences between some full-length and truncated vertebrate MRs in their EC50s for DOC and Swith truncated MRs having higher EC50s (weaker activation) than their corresponding full-length MRs. Moreover, Prog, 19norProg and Spiron, which were transcriptional activators of full-length chicken and zebrafish MRs, were inactive for truncated chicken MR, but retained activity for truncated zebrafish MR. We also find that S, Prog, 19norProg and Spiron compete with Aldo for transcriptional activation of truncated chicken and human MRs, indicating these steroids bind to truncated MRs.We propose that interactions between the A/B/C and D/E domains in vertebrate MRs are important in steroid specificity, with regulation of progestin activation of chicken MR residing in the NTD/DBD and of zebrafish MR in the hinge-LBD. Our data suggests that Prog is a physiological activator of chicken MR, as well as zebrafish and other fish MRs (*11, 18, 30, 34*)

Geller et al. (*17*) found that at 1 nM, Prog, 19norProg and Spiron are transcriptional activators of a Ser810Leu mutant human MR. However, both chicken and zebrafish MRs contain a serine corresponding to Ser810 in wild-type human MR indicating that there are alternative mechanisms for progestin activation of chicken and zebrafish MRs.

## Results

### Comparison of vertebrate MR domains

In Fig. 3, we compare the A/B (NTD), C (DNA-binding domain, DBD), D (hinge region), and E (ligand-binding domain, LBD) domains on human MR with corresponding domains on chicken, alligator, *Xenopus*, and zebrafish MRs. These phylogenetically diverse MRs have strong conservation of the C domain (97-100%) and E domain (77-91%) with substantially less conservation in the A/B domain (36-78%) and D domain (42-78%). 100% identity in the amino acid sequence of the DBD in human, chicken and alligator MRs is important because it eliminates sequence differences in their DBDs as contributing to differences in transcriptional activation by corticosteroids and progestins of these MRs.

### Transcriptional activation by corticosteroids of full-length and truncated human, chicken, alligator, X. laevis, and zebrafish MRs

First, we screened a panel of steroids at 1 nM and 10 nM for transcriptional activation of full-length and truncated human (mammalian), chicken (avian), alligator (reptilian), *Xenopus* (amphibian), and zebrafish (teleost fish) MRs. Aldo, F, B and DOC were similar in activation of full-length human, chicken, alligator and zebrafish MR (Fig. 4, A, C, E, I). Aldo, F and B were strong activators of full length Xenopus MR, while in comparison DOC was a weaker activator (Fig. 4G). S was a good activator, compared to Aldo, of full-length chicken and zebrafish MRs, and a weak activator of full-length human, alligator and Xenopus MRs.

**Figure 4.**
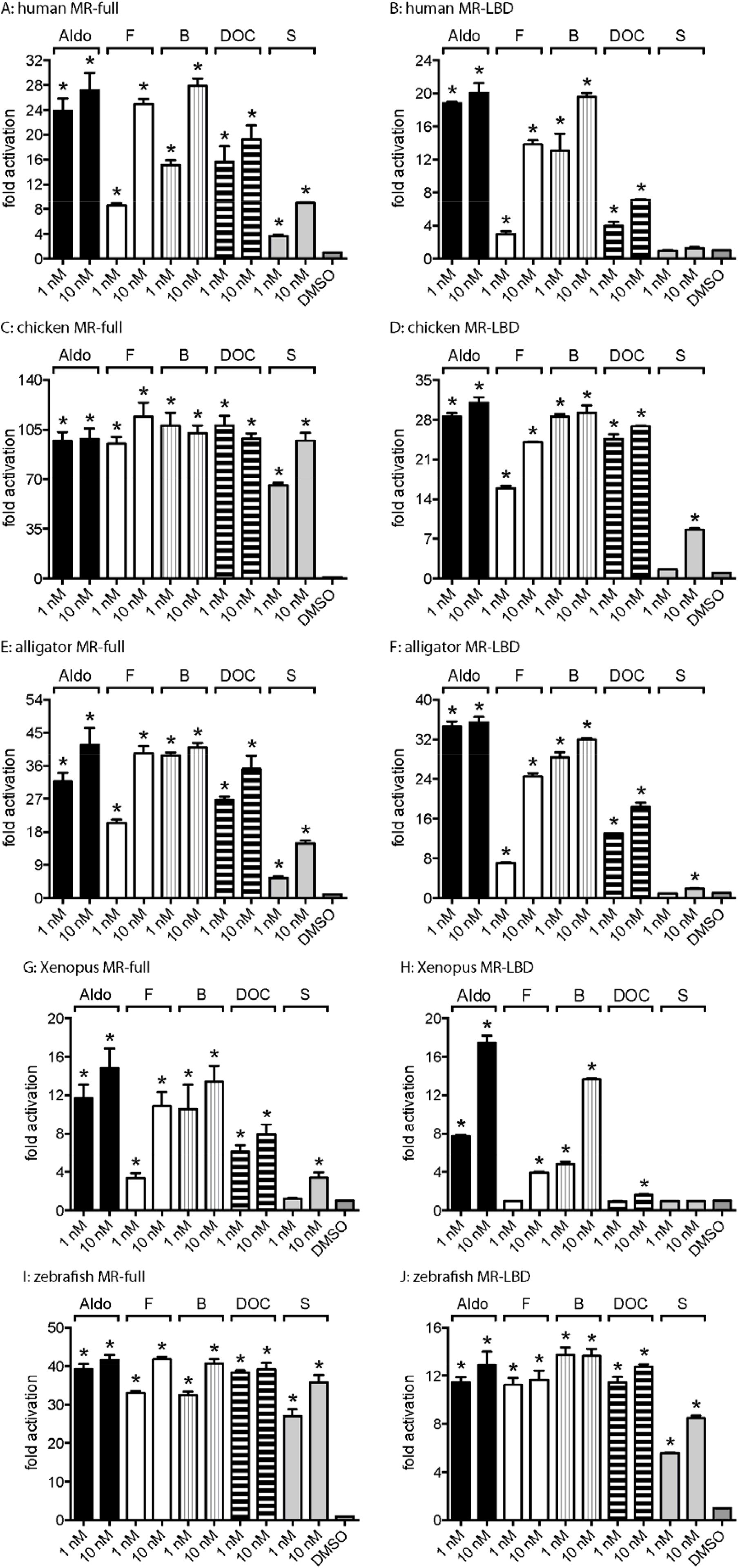
Corticosteroid activation of human, alligator, *Xenopus* and zebrafish full-length MRs and LBD MRs. Full-length human MR (**A**), chicken MR (**C**), alligator MR (**E**), *Xenopus* MR (**G**), and zebrafish MR (**I**) were expressed in HEK293 cells with an MMTV-luciferase reporter. Plasmids for corresponding truncated MRs, human (**B**),chicken (**D**), alligator (**F**), *X. laevis* (**H**) and zebrafish (**J**) containing the D domain and LBD (E domain) fused to a GAL4-DBD were expressed in HEK293 cells with a luciferase reporter. Cells were treated with 1 nM or 10 nM Aldo, F, B, DOC, S or vehicle alone (DMSO). Results are expressed as means ± SEM, n=3.

In contrast, in parallel experiments with truncated MRs, lacking the A/B domain and containing a GAL4-DBD instead of the MR DBD, S had little transcriptional activity at 10 nM for truncated human, chicken,and Xenopus MRs (Fig. 4, B, D, H), in contrast the activity of S for their full-length MRs. S had similar low activity for truncated and full-length alligator MR (Fig. 4F). DOC lost activity, compared to Aldo, for truncated human and chicken MRs and most of its activity for truncated Xenopus MR. However, DOC had similar activity, compared to Aldo, for truncated alligator and zebrafish MRs. At 10 nM, F lost about 25% of its activity, compared to Aldo, for truncated human, chicken and alligator MRs and almost all activity for truncated Xenopus MR (Fig. 4H). The response to corticosteroids by truncated zebrafish MR was different from truncated terrestrial vertebrate MRs (Fig. 4J). Aldo, F, B and DOC had similar activity for truncated and full-length zebrafish MR. At 10 nM, S had about 75% activity for truncated zebrafish MR compared to the full-length zebrafish MR.

### Transcriptional activation by Prog, 19norProg and Spiron of full-length and truncated human, chicken, alligator, X. laevis, and zebrafish MRs

Based on studies showing Prog, 19norProg and Spiron were transcriptional activators of fish MRs (18, 29, 33), we screened these steroids at concentrations of 10 nM, 100 nM and 1 µM for transcriptional activation of full length and truncated vertebrate MRs (Fig. 5). At 1 µM, neither Prog, 19norP nor Spiron were transcriptional activators of full-length human, Xenopus and alligator MRs. However, Prog, 19norP and Spiron activated transcription by full-length zebrafish MR (Fig. 5J) (*30*). Unexpectedly, Prog, 19norP and Spiron activated full-length chicken MR (Fig. 5D). Our finding that Spiron activates full-length chicken MR is in agreement with an earlier report by Proszkowiec-Weglarz and Porter (*61*). Prog, 19norProg and Spiron had no activity for truncated human, alligator and Xenopus MRs (Fig. 5B, F, H). At 1 µM, Prog and 19norProg had about 30% of the activity of 10 nM Aldo for transcriptional activation of truncated chicken MR, while Spiron was inactive. Prog and 19norProg had similar relative activation of truncated zebrafish MR, while spiron had lower activity (Fig. 5J).

**Figure 5.**
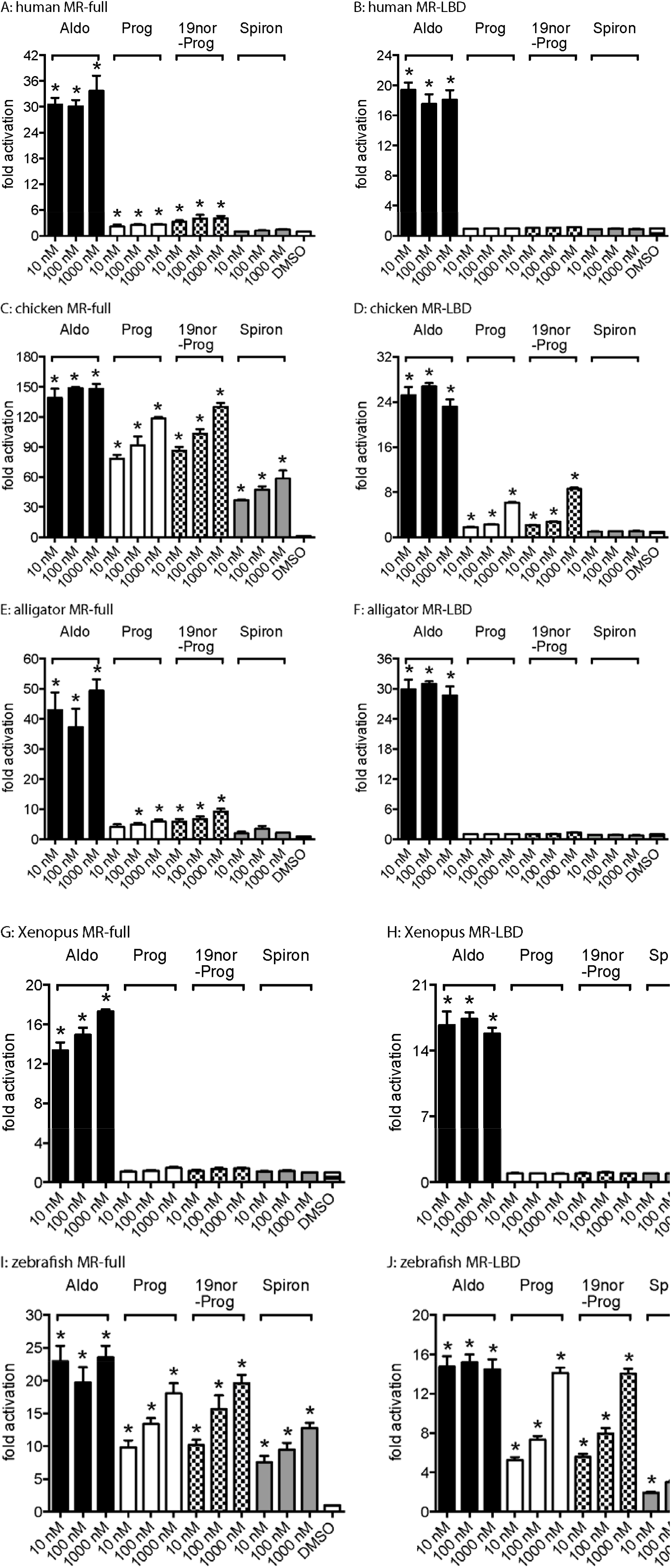
Prog, 19norProg or Spiron activation of human, alligator, *Xenopus* and zebrafish full-length and truncated MRs. Full-length human MR (**A**), chicken MR (**C**), alligator MR (**E**), *Xenopus* MR (**G**), and zebrafish MR (**I**) were expressed in HEK293 cells with an MMTV-luciferase reporter. Plasmids for corresponding truncated MRs, human (**B**),chicken (**D**), alligator (**F**), *X. laevis* (**H**) and zebrafish (**J**) containing the D domain and LBD (E domain) fused to a GAL4-DBD were expressed in HEK293 cells with a luciferase reporter containing GAL4 binding site. Cells were treated with 10 nM, 100 nM or 1 µM Prog, 19norProg or Spiron or vehicle alone (DMSO). Results are expressed as means ± SEM, n=3. Y-axis indicates fold-activation compared to the activity of control vector with vehicle (DMSO) alone as 1. Asterisks indicate significant difference in comparison with DMSO and chemical treatments.

### EC50 values for corticosteroid activation of full-length and truncated human, chicken, alligator, X. laevis and zebrafish MRs. Full-length vertebrate MRs

To gain a quantitative measure of corticosteroid activation of vertebrate MRs, we determined the concentration-dependent activation of full-length vertebrate MRs by Aldo, F, B, DOC and S (Fig. 6, Table 1). Although chicken and zebrafish are phylogenetically distant vertebrates (Fig. 2), full-length chicken and zebrafish MRs have EC50s that are below 1 nM for Aldo, F, B, DOC and S. Human, alligator and Xenopus MRs have strongest responses to Aldo, B and F and weaker responses to DOC and S. Thus, activation of full-length human MR by DOC and S was72% and 42%, respectively, of that of Aldo. In contrast, activation of full-length chicken MR by DOC and S was110% and 112%, respectively, of Aldo, and activation of full-length zebrafish MR by DOC and S was103% and 194%, respectively, of Aldo (Table 1).

**Table 1.** EC50 activities for 3-ketosteroid transcriptional activation of full-length vertebrate MRs.

| MR | Aldo | B | F | DOC | S |
| --- | --- | --- | --- | --- | --- |
|  | EC50 (M) | EC50 (M) | EC50 (M) | EC50 (M) | EC50 (M) |
| Human | $2.7 \times 10^{-10}$ | $1.2 \times 10^{-9}$ | $5.5 \times 10^{-9}$ | $4.2 \times 10^{-10}$ | $3.6 \times 10^{-9}$ |
|  | 100% | 119% | 133% | 74% | 42% |
| Chicken | $6.2 \times 10^{-11}$ | $5.1 \times 10^{-11}$ | $2.8 \times 10^{-10}$ | $3.4 \times 10^{-11}$ | $6.7 \times 10^{-10}$ |
|  | 100% | 109% | 128% | 110% | 112% |
| Alligator | $2.8 \times 10^{-10}$ | $3.6 \times 10^{-10}$ | $6.9 \times 10^{-9}$ | $2.3 \times 10^{-10}$ | $2.7 \times 10^{-9}$ |
|  | 100% | 138% | 176% | 85% | 45% |
| Xenopus | $4.6 \times 10^{-10}$ | $6.2 \times 10^{-10}$ | $1.1 \times 10^{-8}$ | $7.6 \times 10^{-10}$ | $9.1 \times 10^{-9}$ |
|  | 100% | 105% | 126% | 59% | 31% |
| Zebrafish | $8.2 \times 10^{-11}$ | $3.0 \times 10^{-10}$ | $4.4 \times 10^{-10}$ | $6.3 \times 10^{-11}$ | $4.0 \times 10^{-10}$ |
|  | 100% | 112% | 123% | 103% | 94% |

(%) Relative induction compared to Aldosterone induced activation.

**Figure 6.**
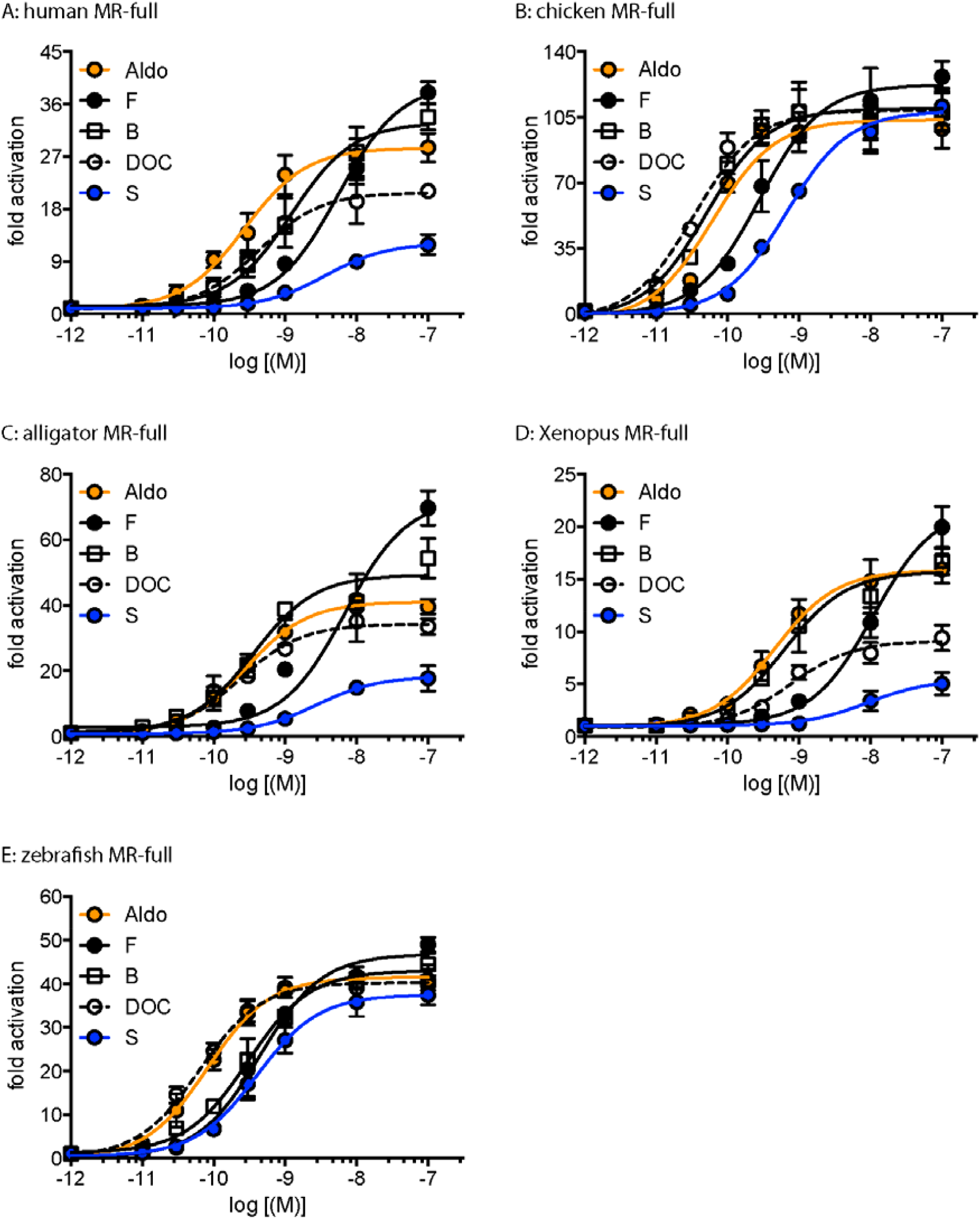
Concentration-dependent transcriptional activation by corticosteroids of full-length human, chicken, alligator, *Xenopus* and zebrafish MRs. Plasmids encoding full-length MRs **A**: human MR, **B**: chicken MR, **C**: alligator MR, **D**: *Xenopus* MR and **E**: zebrafish MR were expressed in HEK293 cells treated with increasing concentrations of steroid or vehicle alone (DMSO). Results are expressed as means ± SEM, n=3. Y-axis indicates fold-activation compared to the activity of control vector with vehicle (DMSO) alone as 1.

### Truncated vertebrate MRs

To investigate the role of the NTD and DBD we determined the concentration-dependent transcriptional activation of truncated terrestrial vertebrate MRs by Aldo, F, B, Aldo, DOC and S. Consistent with our screening studies (Fig. 5), transcriptional activation by S, DOC and F was dramatically lowered for some terrestrial vertebrate MRs that lacked MR NTD- DBD (Fig. 7 and Table 2). For example, S had little activity for truncated human, alligator and Xenopus MRs, and DOC and F had less activity for truncated Xenopus and human MRs. Interestingly, truncated zebrafish MR retained a good response to corticosteroids (Table 2).

**Table 2.** EC50 activities for 3-ketosteroid transcriptional activation of GAL4-DBD-MR-LBD of vertebrate MRs.

| MR | Aldo | B | F | DOC | S |
| --- | --- | --- | --- | --- | --- |
|  | EC50 (M) | EC50 (M) | EC50 (M) | EC50 (M) | EC50 (M) |
| Human | $2.8 \times 10^{-10}$ | $5.9 \times 10^{-10}$ | $3.2 \times 10^{-9}$ | $1.8 \times 10^{-9}$ | - |
|  | 100% | 95% | 74% | 44% | 8% |
| Chicken | $1.3 \times 10^{-10}$ | $1.6 \times 10^{-10}$ | $6.9 \times 10^{-10}$ | $1.7 \times 10^{-10}$ | $4.7 \times 10^{-9}$ |
|  | 100% | 92% | 75% | 92% | 36% |
| Alligator | $3.5 \times 10^{-10}$ | $3.8 \times 10^{-10}$ | $2.3 \times 10^{-9}$ | $5.2 \times 10^{-10}$ | - |
|  | 100% | 88% | 68% | 51% | 8% |
| *Xenopus* | $1.5 \times 10^{-9}$ | $1.9 \times 10^{-9}$ | $1.2 \times 10^{-8}$ | - | - |
|  | 100% | 74% | 37% | 10% | 6% |
| Zebrafish | $2.7 \times 10^{-11}$ | $1.5 \times 10^{-10}$ | $3.1 \times 10^{-10}$ | $1.0 \times 10^{-10}$ | $9.1 \times 10^{-10}$ |
|  | 100% | 96% | 77% | 99% | 67% |

(%) Relative induction compared to Aldosterone induced activation.

**Figure 7.**
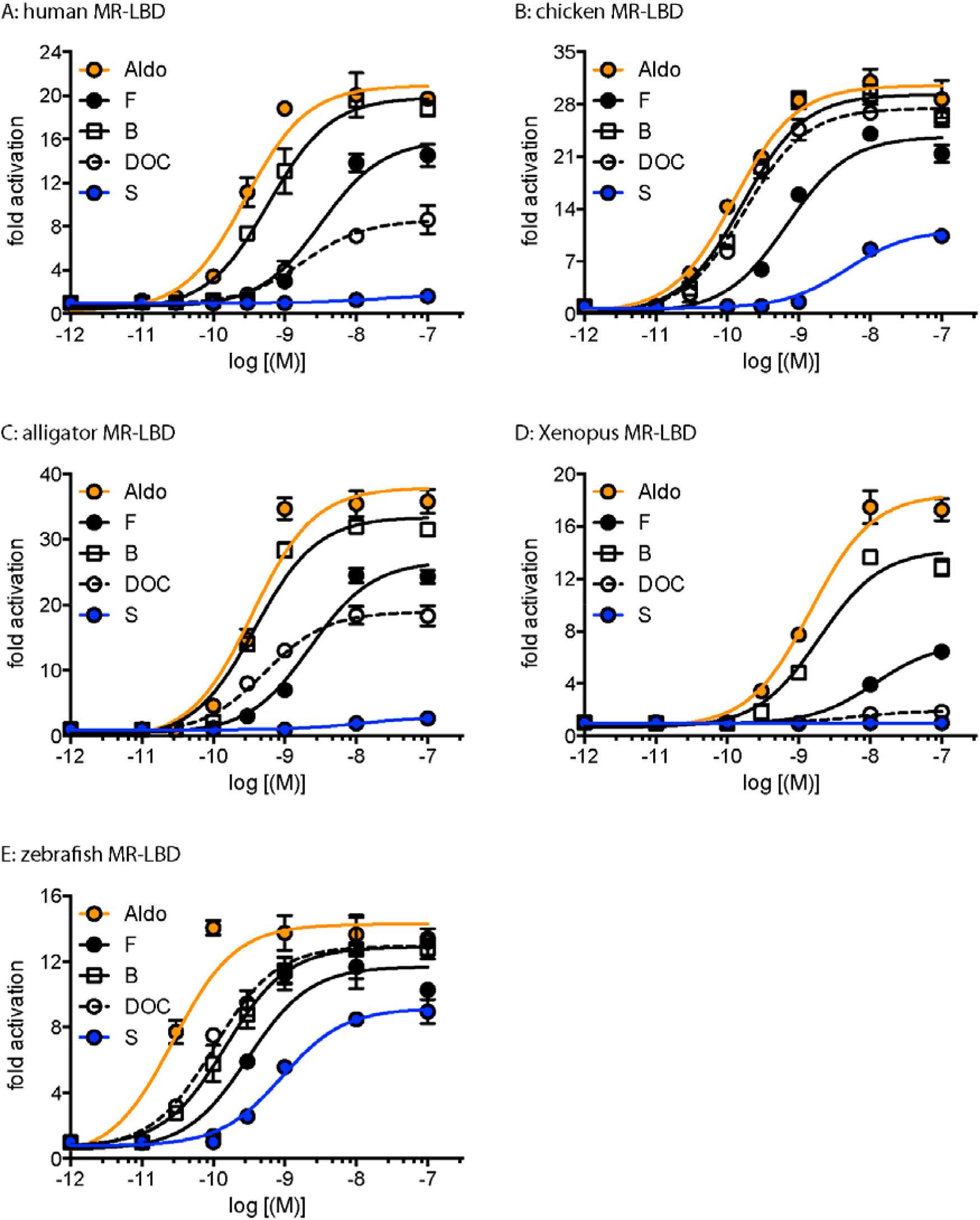
Concentration-dependent transcriptional activation of truncated human, chicken, alligator, *Xenopus* and zebrafish MRs. Plasmids encoding the GAL4-DBD fused to the D domain and LBD of MRs (**A**: human, **B**: chicken, **C**: alligator, **D**: *Xenopus*, **E**: zebrafish)) were expressed in HEK293 cells treated with increasing concentrations of Aldo, F, and DOC or vehicle alone (DMSO). Results are expressed as means ± SEM, n=3. Y-axis indicates fold-activation compared to the activity of control vector with vehicle (DMSO) alone as 1.

### Prog, 19norProg, Spiron, S and DOC inhibit Aldo activation of terrestrial vertebrate MRs

Next, we investigated if Prog, 19norProg, Spiron, S and DOC bind to those truncated MRs that are not activated by these steroids. The effect of these steroids on transcriptional activation by Aldo of truncated human, alligator and Xenopus MRs and of DOC on Aldo activation of Xenopus MR was determined (Fig. 8). All of these steroids inhibited Aldo activation of the truncated MRs indicating that these steroids bind to truncated MRs.

**Figure 8.**
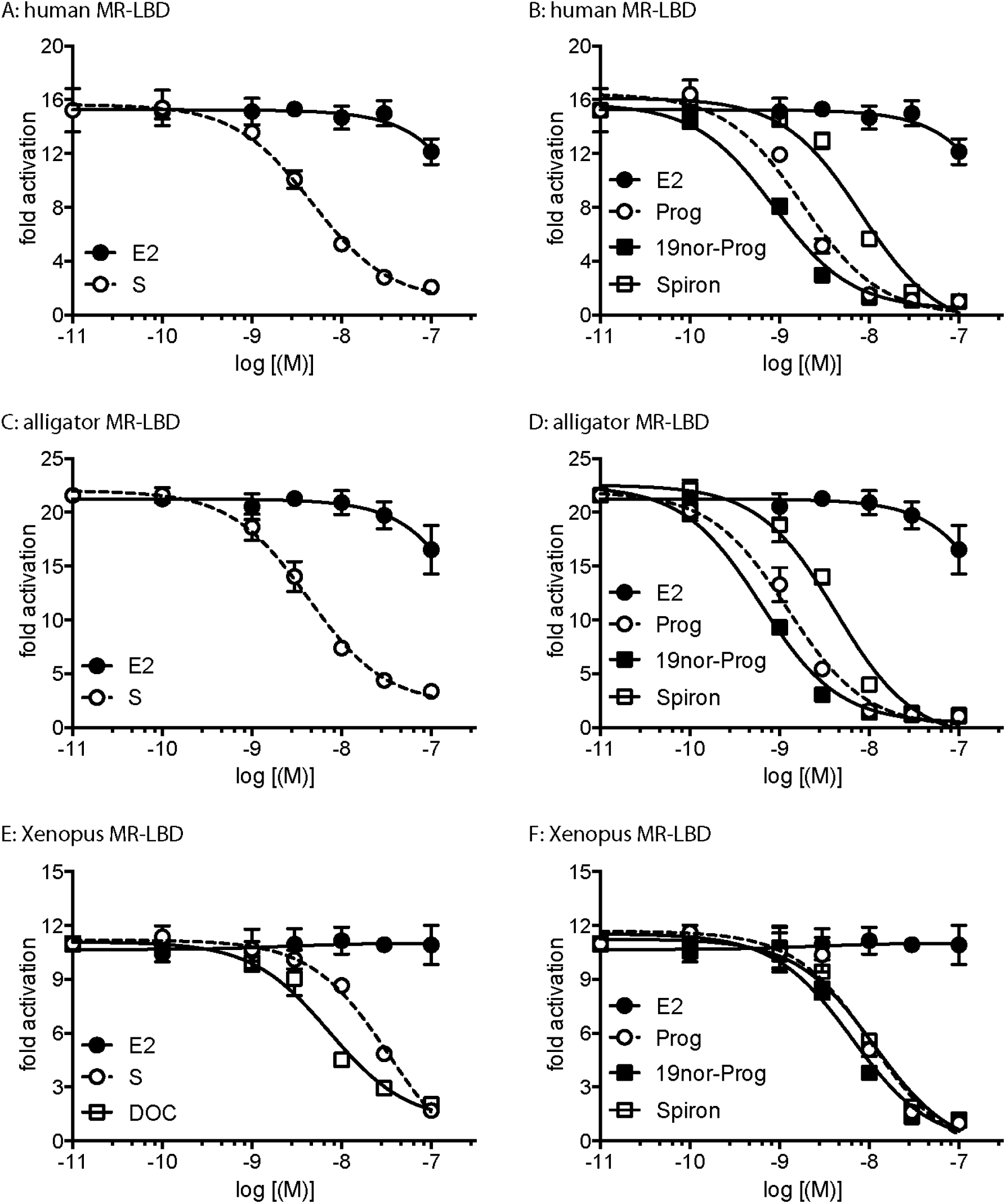
Competition by Prog, 19norProg, Spiron or S for transcriptional activation by Aldo of truncated human, alligator and *Xenopus* MRs. Plasmids encoding the GAL4-DBD fused to the D domain and LBD of MRs (**A, B**: human, **C, D**: alligator, **E, F**: *Xenopus*) were expressed in HEK293 cells. Human and alligator MR were treated with 0.5 nM Aldo and increasing concentrations of Prog, 19norProg, Spiron or S. Xenopus MR was treated with 5 nM Aldo and increasing concentrations of Prog, 19norProg, Spiron, S or DOC. E2 was a control in all experiments. Transcriptional activation of these truncated MRs by Aldo was inhibited by Prog, 19norProg, Spiron, S and DOC indicating that these steroids bind to the truncated MRs. Results are expressed as means ± SEM, n=3. Y-axis indicates fold-activation compared to the activity of control vector with vehicle (DMSO) alone as 1.

## Discussion

Although it is thirty years since the human MR was cloned (10), data on transcriptional activation of vertebrate MRs by corticosteroids and progestins is modest. For the most part, the focus has been on activation of full-length human MR by Aldo and F, and in some studies by B, DOC and S (11, 18, 19, 36, 38, 52, 53). Transcriptional activation by Aldo and B of full-length chicken MR (54) and by Aldo, B, F and DOC full-length alligator MR (55) also has been studied. Data for Prog, 19norProg and Spiron in terrestrial vertebrates is limited to human MR for which Prog and 19norProg have low activity, while Spiron is an MR antagonist (15, 17, 18, 38). In contrast, Prog, 19norProg and Spiron activate the MR in zebrafish, trout, gar and sturgeon (18, 29, 33). Data on the influence of the NTD on transcriptional activation by corticosteroids and progestins is limited to human, rat and zebrafish MRs (19, 29, 36, 56), with no data on chicken, alligator and Xenopus MRs.

Here we fill in some gaps in our knowledge of transcriptional activation by corticosteroids and progestins of full-length and truncated MRs in chicken, alligator and Xenopus and truncated zebrafish MR. The DBDs in human, chicken and alligator MRs are identical and the DBDs in other vertebrate MRs are strongly conserved (Fig. 3) suggesting that differences in transcriptional activation by corticosteroids and progestins of full-length and truncated MRs are mainly are due to interactions between the NTD and hinge-LBD.

The higher EC50s for corticosteroid activation of truncated terrestrial vertebrate MRs compared to full-length MRs (Tables 1 and 2) indicate that the NTD and LBD are important in transcriptional activation of the MR. The loss of activity for truncated MRs varies with the steroid and the vertebrate. Aldo and B have the smallest change in EC50 for full-length and truncated terrestrial vertebrate MRs (Tables 1 and 2). Unexpectedly S has less than 10% of Aldo’s activate for truncated human, alligator and Xenopus MRs, and 36% and 67%, respectively, for chicken and zebrafish MRs, which are the two MRs that are activated by Prog. DOC loses activity for truncated Xenopus MR (10% of Aldo) and has diminished activity for truncated human MR (44% of Aldo). In contrast to terrestrial vertebrate MRs, truncated zebrafish MR retains activity for DOC, S and the other corticosteroids.

Our results are in agreement with Rogerson and Fuller (36) and Pippal et al (19, 29), who found that truncated (GAL4-DBD-MR-LBD) human and zebrafish MRs had lower responses to Aldo than truncated MRs incubated with the NTD domain. They also reported that Aldo could not activate transcription by truncated human Glu962Ala MR, but Aldo could activate NTD+truncated-Glu962Ala MR, demonstrating the importance of human MR NTD in transcriptional activation of human MR. DOC and F promoted an increase in transcriptional activation for NTD+GAL4-DBD-zebrafishMR-LBD, but only had a weak effect for NTD+GAL4-DBD-human LBD. Our results extend this role of the NTD to transcriptional activation of chicken, alligator and Xenopus MRs by Aldo, F, B, DOC and S and transcriptional activation of human MR by B and S (Tables 1 and 2, Fig., 4 and 6).

Prog, 19norProg and Spiron are enigmatic ligands for vertebrate MRs. These steroids are antagonists for human MR (15, 17-19), and as reported here for alligator and Xenopus MRs (Fig. 4), and agonists for zebrafish MR (18, 29), trout MR (33), sturgeon MR and gar MR (18) and as reported here for chicken MR, which was unexpected (Fig. 4). Chicken MR also is unusual because co-expression of chicken MR and GR does not affect their transcriptional activity (*61*), which differs from the clear effect on transcription of co-expression of human MR and GR (*62-64*) and trout MR and GR (*63*).

Prog may be a transcriptional activator of chicken MR in tissues that contain 11β-HSD2, which converts B to inactive 11-dehydrocorticosterone (*5, 8, 22, 24*). Due to the absence of an 11β-hydroxyl on Prog, it is inert to 11β-HSD2, which would convert about 90% of B to 11-dehydrocorticosterone (*14, 65*). The concentrations of Prog and B in chicken blood are 4.7 nM and 15.3 nM, respectively (*66*), and an upper bound for Aldo in chicken blood is 0.4 nM (*67*). Thus, the physiological concentration of Prog is sufficient to occupy chicken MR.

For terrestrial vertebrate MRs, the only previous example of a Prog-activated MR was the Ser810Leu mutant human MR, which was studied by Geller et al. (17). Ser810Leu MR was activated by 1 nM Prog, 19norProg and Spiron. However, both chicken and zebrafish MRs, as well as other Prog-activated fish MRs (Figure 2) (*18, 30, 34*), contain a serine corresponding to Ser810 in wild-type human MR. This indicates that alternative mechanisms are involved in progestin activation of chicken, zebrafish and other fish MRs. Our experiments with truncated chicken and zebrafish MRs provide clues to regulation of Prog activation of chicken and zebrafish MRs. The lack of Prog, 19norProg and Spiron activation of truncated chicken MR indicates that allosteric interactions between the NTD and hinge-LBD domains of full-length chicken MR are important in transcriptional activation by these steroids, while activation of truncated zebrafish MR by Prog, 19norProg and Spiron indicates that the hinge-LBD domains are important in transcriptional activation of full-length zebrafish MR.

## Materials and Methods

### Chemical reagents

Steroids with their CAS numbers: cortisol (F): 50-23-7, corticosterone (B): 50-22-6, aldosterone (Aldo): 52-39-1, 11-deoxycorticosterone (DOC): 64-85-7, 11-deoxycortisol (S): 152-58-9, progesterone (Prog): 57-83-0, 19nor-progesterone (19norProg): 472-54-8 and spironolactone (Spiron): 57-01-7 were purchased from Sigma-Aldrich. For reporter gene assays, all hormones were dissolved in dimethylsulfoxide (DMSO) and the final concentration of DMSO in the culture medium did not exceed 0.1% v/v, which was not toxic.

### Construction of plasmid vectors

The full-coding regions and D/E domains of the MR from human, chicken, alligator, frog (Xenopus) and zebrafish were amplified by PCR with KOD DNA polymerase. The PCR products were gel-purified and ligated into pcDNA3.1 vector (KpnI-NotI site for human MR, and BamHI-NotI site for chicken, alligator, frog and zebrafish MRs) for the full-coding region or pBIND vector (MluI-NotI site for human, chicken, frog and zebrafish MR, and MluI-KpnI site for alligator MR) for D-E domains. As shown in Fig. 3, the D domain begins at human MR (732), chicken MR (729), alligator MR (733), frog MR (727), and zebrafish MR (718).

### Transactivation Assay and Statistical Methods

Human embryonic kidney 293 (Hek293) cells were used in the reporter gene assay, and transfection and reporter assays were carried out as described previously (18, 51). All transfections were performed at least three times, employing triplicate sample points in each experiment. The values shown are mean ± SEM from three separate experiments, and dose-response data and EC50 were analyzed using GraphPad Prism. One–way ANOVA followed by Dunnett’s multiple comparisons test was performed for statistical analyses. Differences were considered statistically signify at *P <* 0.05.

## Acknowledgments

**Funding**: K.O. was supported by the Japan Society for the Promotion of Science (JSPS) Research Fellowships for Young Scientists. This work was supported in part by Grants-in-Aid for Scientific Research 23570067 and 26440159 (YK) from the Ministry of Education, Culture, Sports, Science and Technology of Japan. M.E.B. was supported by Research fund #3096.

## Author contributions

Y.K. and K.O. carried out the research. Y.K. and M.E.B. conceived and designed the experiments and wrote the paper. All authors gave final approval for publication.

## References

1 G. V. Markov et al., Independent elaboration of steroid hormone signaling pathways in metazoans. Proc Natl Acad Sci U S A 106, 11913–11918 (2009).

2 J. T. Bridgham et al., Protein evolution by molecular tinkering: diversification of the nuclear receptor superfamily from a ligand-dependent ancestor. PLoS biology 8, (2010).

3 S. Bertrand, M. R. Belgacem, H. Escriva, Nuclear hormone receptors in chordates. Molecular and cellular endocrinology 334, 67–75 (2011).

4 M. E. Baker, D. R. Nelson, R. A. Studer, Origin of the response to adrenal and sex steroids: Roles of promiscuity and co-evolution of enzymes and steroid receptors. J Steroid Biochem Mol Biol 151, 12–24 (2015).

5 J. W. Funder, Aldosterone and mineralocorticoid receptors: a personal reflection. Molecular and cellular endocrinology 350, 146–150 (2012).

6 U. A. Hawkins, E. P. Gomez-Sanchez, C. M. Gomez-Sanchez, C. E. Gomez-Sanchez, The ubiquitous mineralocorticoid receptor: clinical implications. Curr Hypertens Rep 14, 573–580 (2012).

7 L. Martinerie et al., The mineralocorticoid signaling pathway throughout development: expression, regulation and pathophysiological implications. Biochimie 95, 148–157 (2013).

8 B. C. Rossier, M. E. Baker, R. A. Studer, Epithelial sodium transport and its control by aldosterone: the story of our internal environment revisited. Physiological reviews 95, 297–340 (2015).

9 F. Jaisser, N. Farman, Emerging Roles of the Mineralocorticoid Receptor in Pathology: Toward New Paradigms in Clinical Pharmacology. Pharmacological reviews 68, 49–75 (2016).

10 J. L. Arriza et al., Cloning of human mineralocorticoid receptor complementary DNA: structural and functional kinship with the glucocorticoid receptor. Science 237, 268–275 (1987).

11 M. E. Baker, Y. Katsu, 30 YEARS OF THE MINERALOCORTICOID RECEPTOR: Evolution of the mineralocorticoid receptor: sequence, structure and function. J Endocrinol 234, T1–T16 (2017).

12 R. Rupprecht et al., Pharmacological and functional characterization of human mineralocorticoid and glucocorticoid receptor ligands. Eur J Pharmacol 247, 145–154 (1993).

13 Z. S. Krozowski, J. W. Funder, Renal mineralocorticoid receptors and hippocampal corticosterone-binding species have identical intrinsic steroid specificity. Proc Natl Acad Sci U S A 80, 6056–6060 (1983).

14 K. Myles, J. W. Funder, Progesterone binding to mineralocorticoid receptors: in vitro and in vivo studies. Am J Physiol 270, E601–607 (1996).

15 K. Myles, J. W. Funder, Type I (mineralocorticoid) receptors in the guinea pig. Am J Physiol 267, E268–272 (1994).

16 C. Hellal-Levy et al., Specific hydroxylations determine selective corticosteroid recognition by human glucocorticoid and mineralocorticoid receptors. FEBS Lett 464, 9–13 (1999).

17 D. S. Geller et al., Activating mineralocorticoid receptor mutation in hypertension exacerbated by pregnancy. Science 289, 119–123 (2000).

18 A. Sugimoto et al., Corticosteroid and progesterone transactivation of mineralocorticoid receptors from Amur sturgeon and tropical gar. Biochem J 473, 3655–3665 (2016).

19 J. B. Pippal, Y. Yao, F. M. Rogerson, P. J. Fuller, Structural and functional characterization of the interdomain interaction in the mineralocorticoid receptor. Mol Endocrinol 23, 1360–1370 (2009).

20 M. E. Baker, J. W. Funder, S. R. Kattoula, Evolution of hormone selectivity in glucocorticoid and mineralocorticoid receptors. J Steroid Biochem Mol Biol 137, 57–70 (2013).

21 J. Funder, K. Myles, Exclusion of corticosterone from epithelial mineralocorticoid receptors is insufficient for selectivity of aldosterone action: in vivo binding studies. Endocrinology 137, 5264–5268 (1996).

22 E. Gomez-Sanchez, C. E. Gomez-Sanchez, The multifaceted mineralocorticoid receptor. Compr Physiol 4, 965–994 (2014).

23 A. Odermatt, A. G. Atanasov, Mineralocorticoid receptors: emerging complexity and functional diversity. Steroids 74, 163–171 (2009).

24 A. Odermatt, D. V. Kratschmar, Tissue-specific modulation of mineralocorticoid receptor function by 11beta-hydroxysteroid dehydrogenases: an overview. Molecular and cellular endocrinology 350, 168–186 (2012).

25 K. Chapman, M. Holmes, J. Seckl, 11beta-hydroxysteroid dehydrogenases: intracellular gate-keepers of tissue glucocorticoid action. Physiological reviews 93, 1139–1206 (2013).

26 E. P. Gomez-Sanchez, Mineralocorticoid receptors in the brain and cardiovascular regulation: minority rule? Trends Endocrinol Metab 22, 179–187 (2011).

27 J. W. Funder, Aldosterone and mineralocorticoid receptors in the cardiovascular system. Prog Cardiovasc Dis 52, 393–400 (2010).

28 J. L. Arriza, R. B. Simerly, L. W. Swanson, R. M. Evans, The neuronal mineralocorticoid receptor as a mediator of glucocorticoid response. Neuron 1, 887–900 (1988).

29 R. Rupprecht et al., Transactivation and synergistic properties of the mineralocorticoid receptor: relationship to the glucocorticoid receptor. Molecular endocrinology (Baltimore, Md) 7, 597–603 (1993).

30 J. B. Pippal, C. M. Cheung, Y. Z. Yao, F. E. Brennan, P. J. Fuller, Characterization of the zebrafish (Danio rerio) mineralocorticoid receptor. Molecular and cellular endocrinology 332, 58–66 (2011).

31 A. S. Arterbery et al., Evolution of ligand specificity in vertebrate corticosteroid receptors. BMC Evol Biol 11, 14 (2011).

32 A. K. Greenwood et al., Multiple corticosteroid receptors in a teleost fish: distinct sequences, expression patterns, and transcriptional activities. Endocrinology 144, 4226–4236 (2003).

33 E. H. Stolte et al., Corticosteroid receptors involved in stress regulation in common carp, Cyprinus carpio. J Endocrinol 198, 403–417 (2008).

34 A. Sturm et al., 11-deoxycorticosterone is a potent agonist of the rainbow trout (Oncorhynchus mykiss) mineralocorticoid receptor. Endocrinology 146, 47–55 (2005).

35 P. Huang, V. Chandra, F. Rastinejad, Structural overview of the nuclear receptor superfamily: insights into physiology and therapeutics. Annu Rev Physiol 72, 247–272 (2010).

36 F. Rastinejad, P. Huang, V. Chandra, S. Khorasanizadeh, Understanding nuclear receptor form and function using structural biology. Journal of molecular endocrinology 51, T1–T21 (2013).

37 F. M. Rogerson, P. J. Fuller, Interdomain interactions in the mineralocorticoid receptor. Molecular and cellular endocrinology 200, 45–55 (2003).

38 Y. Li, K. Suino, J. Daugherty, H. E. Xu, Structural and biochemical mechanisms for the specificity of hormone binding and coactivator assembly by mineralocorticoid receptor. Mol Cell 19, 367–380 (2005).

39 R. K. Bledsoe et al., A ligand-mediated hydrogen bond network required for the activation of the mineralocorticoid receptor. The Journal of biological chemistry 280, 31283–31293 (2005).

40 K. Edman et al., Ligand Binding Mechanism in Steroid Receptors: From Conserved Plasticity to Differential Evolutionary Constraints. Structure 23, 2280–2290 (2015).

41 J. Fagart et al., Crystal structure of a mutant mineralocorticoid receptor responsible for hypertension. Nature structural & molecular biology 12, 554–555 (2005).

42 P. J. Fuller, Novel interactions of the mineralocorticoid receptor. Molecular and cellular endocrinology 408, 33–37 (2015).

43 Z. X. Zhou, M. V. Lane, J. A. Kemppainen, F. S. French, E. M. Wilson, Specificity of ligand-dependent androgen receptor stabilization: receptor domain interactions influence ligand dissociation and receptor stability. Mol Endocrinol 9, 208–218 (1995).

44 M. J. Tetel, P. H. Giangrande, S. A. Leonhardt, D. P. McDonnell, D. P. Edwards, Hormone-dependent interaction between the amino- and carboxyl-terminal domains of progesterone receptor in vitro and in vivo. Mol Endocrinol 13, 910–924 (1999).

45 R. Metivier, G. Penot, G. Flouriot, F. Pakdel, Synergism between ERalpha transactivation function 1 (AF-1) and AF-2 mediated by steroid receptor coactivator protein-1: requirement for the AF-1 alpha-helical core and for a direct interaction between the N- and C-terminal domains. Mol Endocrinol 15, 1953–1970 (2001).

46 J. Thompson, F. Saatcioglu, O. A. Janne, J. J. Palvimo, Disrupted amino- and carboxyl-terminal interactions of the androgen receptor are linked to androgen insensitivity. Mol Endocrinol 15, 923–935 (2001).

47 R. Kumar, G. Litwack, Structural and functional relationships of the steroid hormone receptors’ N-terminal transactivation domain. Steroids 74, 877–883 (2009).

48 K. Fischer, S. M. Kelly, K. Watt, N. C. Price, I. J. McEwan, Conformation of the mineralocorticoid receptor N-terminal domain: evidence for induced and stable structure. Mol Endocrinol 24, 1935–1948 (2010).

49 S. S. Simons, Jr., R. Kumar, Variable steroid receptor responses: Intrinsically disordered AF1 is the key. Molecular and cellular endocrinology 376, 81–84 (2013).

50 A. Christopoulos et al., International union of basic and clinical pharmacology. XC. multisite pharmacology: recommendations for the nomenclature of receptor allosterism and allosteric ligands. Pharmacological reviews 66, 918–947 (2014).

51 K. Oka et al., Allosteric role of the amino-terminal A/B domain on corticosteroid transactivation of gar and human glucocorticoid receptors. J Steroid Biochem Mol Biol 154, 112–119 (2015).

52 Y. Katsu, S. Kohno, K. Oka, M. E. Baker, Evolution of corticosteroid specificity for human, chicken, alligator and frog glucocorticoid receptors. Steroids 113, 38–45 (2016).

53 N. R. Bury, A. Sturm, Evolution of the corticosteroid receptor signalling pathway in fish. Gen Comp Endocrinol 153, 47–56 (2007).

54 T. Sakamoto et al., Corticosteroids stimulate the amphibious behavior in mudskipper: potential role of mineralocorticoid receptors in teleost fish. Physiol Behav 104, 923–928 (2011).

55 H. Takahashi, T. Sakamoto, The role of ‘mineralocorticoids’ in teleost fish: relative importance of glucocorticoid signaling in the osmoregulation and ‘central’ actions of mineralocorticoid receptor. Gen Comp Endocrinol 181, 223–228 (2013).

56 J. Q. Jiang, G. Young, T. Kobayashi, Y. Nagahama, Eel (Anguilla japonica) testis 11beta-hydroxylase gene is expressed in interrenal tissue and its product lacks aldosterone synthesizing activity. Molecular and cellular endocrinology 146, 207–211 (1998).

57 D. A. Close, S. S. Yun, S. D. McCormick, A. J. Wildbill, W. Li, 11-deoxycortisol is a corticosteroid hormone in the lamprey. Proc Natl Acad Sci U S A 107, 13942–13947 (2010).

58 B. W. Roberts et al., Regulation of a putative corticosteroid, 17,21-dihydroxypregn-4-ene,3,20-one, in sea lamprey, Petromyzon marinus. Gen Comp Endocrinol 196, 17–25 (2014).

59 N. Saitou, M. Nei, The neighbor-joining method: a new method for reconstructing phylogenetic trees. Molecular biology and evolution 4, 406–425 (1987).

60 J. D. Thompson, D. G. Higgins, T. J. Gibson, CLUSTAL W: improving the sensitivity of progressive multiple sequence alignment through sequence weighting, position-specific gap penalties and weight matrix choice. Nucleic acids research 22, 4673–4680 (1994).

61 M. Proszkowiec-Weglarz, T. E. Porter, Functional characterization of chicken glucocorticoid and mineralocorticoid receptors. American journal of physiology. Regulatory, integrative and comparative physiology 298, R1257–1268 (2010).

62 W. Liu, J. Wang, N. K. Sauter, D. Pearce, Steroid receptor heterodimerization demonstrated in vitro and in vivo. Proc Natl Acad Sci U S A 92, 12480–12484 (1995).

63 P. Kiilerich et al., Interaction between the trout mineralocorticoid and glucocorticoid receptors in vitro. J Mol Endocrinol 55, 55–68 (2015).

64 X. M. Ou, J. M. Storring, N. Kushwaha, P. R. Albert, Heterodimerization of mineralocorticoid and glucocorticoid receptors at a novel negative response element of the 5-HT1A receptor gene. The Journal of biological chemistry 276, 14299–14307 (2001).

65 J. W. Funder, Mineralocorticoid receptor activation and specificity-conferring mechanisms: a brief history. Journal of Endocrinology 234, T17–T21 (2017).

66 M. D. Taves et al., Locally elevated cortisol in lymphoid organs of the developing zebra finch but not Japanese quail or chicken. Dev Comp Immunol 54, 116–125 (2016).

67 T. E. Porter, S. Ghavam, M. Muchow, I. Bossis, L. Ellestad, Cloning of partial cDNAs for the chicken glucocorticoid and mineralocorticoid receptors and characterization of mRNA levels in the anterior pituitary gland during chick embryonic development. Domestic animal endocrinology 33, 226–239 (2007).

